# Differential beta and gamma activity modulation during unimanual and bimanual motor learning

**DOI:** 10.1101/2024.11.11.623009

**Authors:** Min Wu, Marleen J. Schoenfeld, Carl Lindersson, Sven Braeutigam, Catharina Zich, Charlotte J. Stagg

## Abstract

Movement-related dynamics in the beta and gamma bands have been studied in relation to motor execution and learning during unimanual movements, but their roles in complex bimanual tasks remain largely unexplored. This study aimed to investigate how beta and gamma activity differs between unimanual and bimanual movements, and how these neural signatures evolve during the learning process. Our motor task incorporated varying levels of bimanual interaction: unimanual, bimanual-equal, and bimanual-unequal. Magnetoencephalography data were recorded during task performance, and beta and gamma dynamics were quantified. As expected, increasing task complexity from unimanual to bimanual-equal, and then to bimanual-unequal movements resulted in slower and less accurate performance. Across all conditions, significant beta event-related desynchronization (ERD) and gamma event-related synchronization (ERS) were observed during movement, as well as beta ERS after movement. Bimanual movements exhibited greater beta ERD, beta ERS, and gamma ERS compared to unimanual movements. With practice, participants demonstrated faster and more accurate movements, accompanied by enhanced beta ERS responses. Furthermore, learning-related reductions in errors correlated with increases in beta ERS. These findings suggest the distinct behavioural and neural demands of unimanual versus bimanual movements and highlight the important role of beta dynamics in motor performance and learning.

## 1. Introduction

Complex bimanual movements requiring a finely tuned coordination between the upper limbs are essential for daily activities. However, despite their ubiquity, research into the neural mechanisms underlying motor function has predominantly focused on unimanual motor tasks (Alhussein and Smith, 2021; Dekleva et al., 2018; Gonzalez Castro et al., 2014), or simple bimanual motor tasks involving only finger actions, such as finger tapping or figure oscillation (Brandes et al., 2017; Liuzzi et al., 2011; Mechsner et al., 2001). This focus extends to research on motor skill acquisition, which has traditionally prioritized unimanual movements over more complex bimanual movements (Andres and Gerloff, 1999). However, investigating complex bimanual movements that more accurately reflect real-life motor function reveals rich insights into the neural mechanisms of motor control and learning.

Behavioural and neurophysiological studies have revealed important distinctions between unimanual and bimanual movements, as well as differences between bimanual tasks. Unimanual movements are generally performed with greater precision and ease than bimanual movements which inherently depend on the coordination between both hands, resulting in longer movement times and larger errors (Sisti et al., 2011). Moreover, bimanual movements require greater interhemispheric cooperation, with efficient communication between the motor cortices becoming increasingly crucial as task complexity increases (Rueda-Delgado et al., 2017; Serrien, 2008).

Brain activity in the beta band (13-30 Hz) plays a central role in movement, showing a significant decrease in power during movement execution [the event-related desynchronization (ERD)], and an increase in power following movement cessation [the event-related synchronization (ERS)] (Pfurtscheller et al., 1997b; Pfurtscheller and Aranibar, 1977). The beta ERD is hypothesised to be related to task complexity: bimanual tasks show stronger beta ERD compared to unimanual tasks, with anti-phase bimanual tasks exhibiting more significant beta ERD than in-phase tasks, and complex, role-differentiated tasks leading to more pronounced beta ERD than simpler, non-role-differentiated tasks (Gross et al., 2005; Rudisch et al., 2023). However, some studies report contradictory findings (e.g. Houweling et al., 2010, 2008), highlighting the complex role of beta activity during different motor tasks.

Movement-related beta activity is also postulated to have a role in acquisition of motor skills. Learning-related increases in beta ERS are evident, with stronger beta ERS relating to higher movement accuracy in unimanual tasks (Tan et al., 2014b). Additionally, in simple reaching tasks a practice-related strengthening of the beta ERD, the beta ERS, and the modulation depth (i.e., the difference between ERD and ERS) has been observed (Nelson et al., 2017; Ricci et al., 2019). However, these studies, primarily focused on unimanual movements limiting the generalization of their results to more complex bimanual movements, which constitute most human motor activities in daily life.

There is also a growing interest in the role of mid-gamma activity (60-90 Hz) in motor control and motor learning (Nowak et al., 2018; Zich et al., 2021). Gamma is a prokinetic emergent property of the motor cortices, and increases in power during motor execution – the gamma ERS (Crone et al., 1998), and local gamma power has been linked to motor performance and motor learning in unimanual tasks (Zich et al., 2021). Conversely, a recent study demonstrated a decrease in gamma ERS after practice, suggesting a potential shift in attentional demands rather than in movement characteristics (Tatti et al., 2023). However, the role of gamma dynamics in bimanual movements, particularly in bimanual movements of varying complexities, remains largely unexplored.

This study was therefore designed with two primary objectives: (1) to investigate beta and gamma dynamics across unimanual and diverse bimanual movements, and (2) to examine how these neural signatures change throughout the learning process. To achieve these goals, we used a motor task we recently developed (Schoenfeld et al., 2021), where participants were required to move a cursor along an angled path, with each hand controlling movement along orthogonal axes. This approach allowed us to assess different levels of bimanual interaction by incorporating paths that required either a unimanual response, an equal contribution from both hands (bimanual equal), or differing contributions from each hand (bimanual unequal). We investigated beta ERD and gamma ERS during movement, and beta ERS after movement across different conditions, and tracked how these dynamics changed with practice. We hypothesised that these dynamics would differ between movement conditions and would be significantly modulated by practice. Specifically, in line with our previous work, we hypothesised that beta ERD would be more pronounced during learning, and correlate with changes in movement time, and beta ERS would increase during learning and correlate with changes in accuracy.

## 2. Methods

### 2.1 Participants

Forty-three healthy participants (27 females; mean age ± SD = 24.1 ± 4.2 years) were recruited for this study. All participants were right-handed according to the Edinburgh Handedness Inventory (Oldfield, 1971) and had normal or corrected-to-normal vision. All fulfilled the following inclusion criteria: (1) no history of neurological or psychiatric disease; (2) no physical disability of the arms or wrists; and (3) no use of drugs affecting the central nervous system. The study was approved by the Central University Research Ethics Committee, University of Oxford (MSD-IDREC-R68649/RE002). All subjects provided written informed consent in accordance with the Declaration of Helsinki.

### 2.2 Experimental paradigm

Participants were seated upright in the MEGIN TRIUX neo chair with their heads positioned within the magnetoencephalography (MEG) sensor array. Before the bimanual motor learning task, the force grippers were calibrated for each participant. This calibration required participants to exert maximum effort by squeezing each gripper five times, which established the maximum force for each gripper individually by using the peak maximum relative to the gripper’s system offset. During the motor learning task, participants were first familiarised with the task using one orientation street. This was followed by 50 bimanual motor training trials. Each trial consisted of six streets: two unimanual streets at 0° and 90°, two bimanual-equal streets at 45°, and two bimanual-unequal streets at 22.5° and 67.5° (detailed below). Each street started with a 5-second fixation cross at the centre of the screen. Following this, a street was displayed whereby the angle of the street indicated the required cursor movement. Participants then started navigating the cursor from the start position (bottom-centre of the street) along the ideal path in the middle of the street. If the cursor contacted either side of the street, the cursor was reset to the starting position. Upon reaching the end of the street, there was a 5-second resting interval before the next street was presented. This procedure was repeated for each of the six streets. After each trial feedback on the participant’s performance, i.e., movement time (MT) and accuracy, for that trial was visually presented.

Participants were instructed to control the cursor using the left hand for vertical movements and the right hand for horizontal movements. Each street required specific hand contributions based on its angle (Figure 1C). In the unimanual condition, the vertical (0°) streets were controlled entirely by the left hand, and the horizontal (90°) streets entirely by the right hand, effectively resulting in hand contributions of 100% from one hand and 0% from the other. In the bimanual-equal condition, the 45° streets required an equal contribution of 50% effort from each hand. In the bimanual-unequal condition, the streets angled at 22.5° and 67.5° required asymmetric contributions, with the 22.5° streets requiring 75% effort from the left hand and 25% from the right, and the 67.5° streets requiring 25% effort from the left and 75% from the right (Figure 1A-C). To prevent memorisation of the order of the streets, the order of the six streets was pseudo-randomised across trials, whereby two unimanual streets or two bimanual-equal streets never occurred successively.

**Figure 1.**
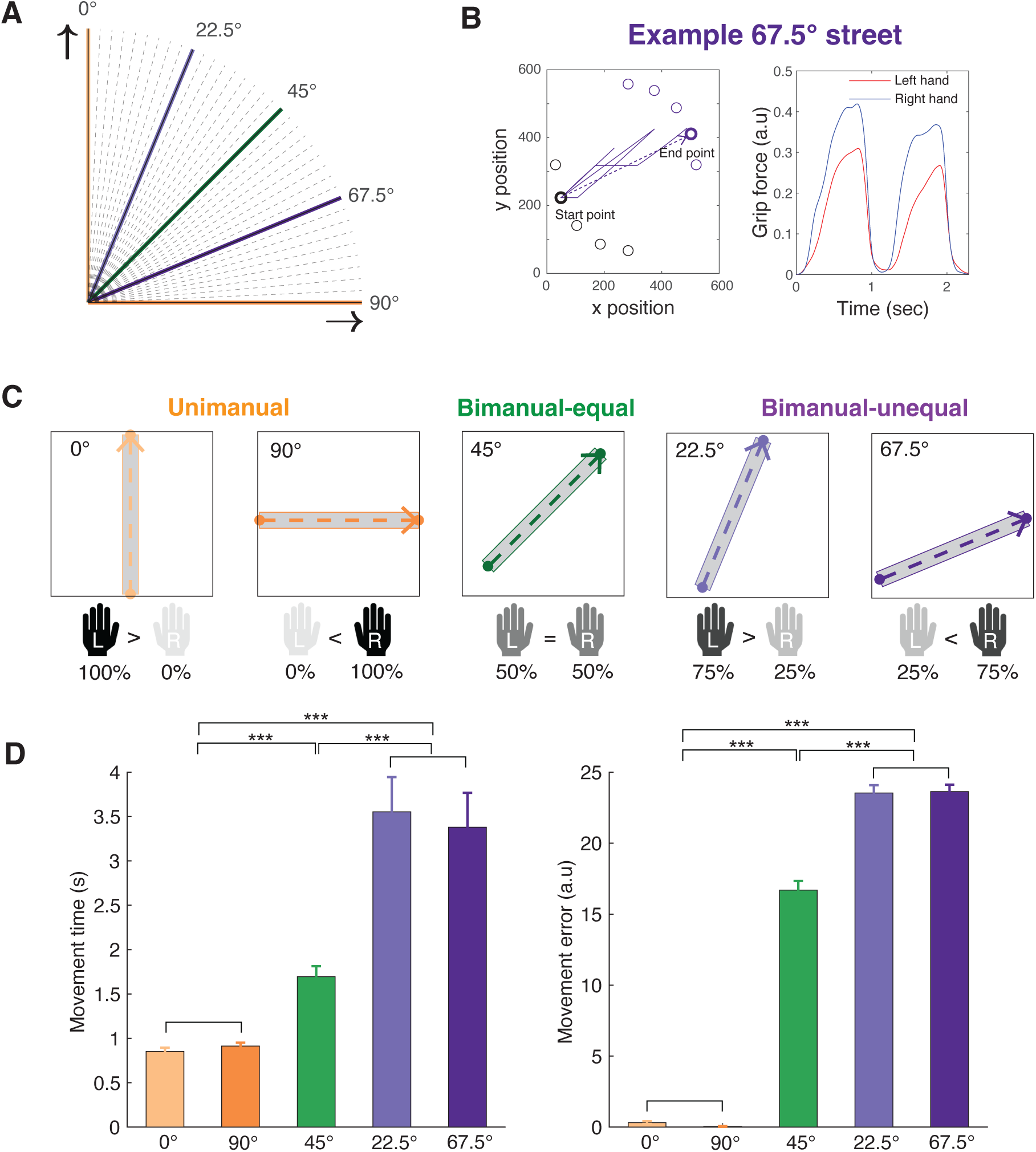
Design of the motor task and behavioural performance. (A) Streets are defined at angles of 0°, 22.5°, 45°, 67.5°, and 90° to create varying degrees of bimanual hand contribution. The unimanual condition is represented in orange (0° and 90°), the bimanual-equal condition in green (45°), and the bimanual-unequal condition in purple (22.5° and 67.5°). (B) Example cursor movement path and hand contribution from a single 67.5° street. The purple lines depict the actual cursor movement path, and the dashed purple line represents the ideal path (left). In this instance, the right hand exerts greater force than the left hand to control the cursor along the 67.5° street (right). (C) Hand contribution for streets of different angles. Each hand controls movement along a different axis on the screen: the left hand controls upward movement, and the right hand controls rightward movement. Different angles correspond to specific hand contributions. In the unimanual condition (0° and 90° streets), only one hand is used (left and right hand, respectively). In the bimanual-equal condition (45° street), both hands contribute equally (50%:50%). In the bimanual-unequal condition (22.5° and 67.5° streets), unequal contribution (75%:25% and 25%:75%, respectively) is required. (D) Behavioural performance for unimanual, bimanual-equal, and bimanual-unequal conditions. Movement time and error are smaller in unimanual conditions than in bimanual-equal, and smaller in bimanual-equal than in bimanual-unequal conditions. Error bars represent the standard error of the mean (S.E.M.). Asterisks indicate significance (*** *p* < 0.001).

Cursor movement required at least 15% of the participant’s maximum force. The participant’s maximum force moved the cursor 25 pixels per refresh rate of the monitor (60 Hz). Once the cursor reached the end of a street, participants needed to reduce their grip to less than 2% of their maximum force to proceed to the fixation cross. This task design built upon our previous work (Schoenfeld et al., 2021), with the primary modification being an extended time between streets allowing for a post-movement beta ERS and a baseline period for the next street.

### 2.3 Behavioural data analysis

Movement time and error were calculated for each street. Movement time was defined as the time between movement onset and movement offset. Movement onset was defined as the time when the gripper force exceeded the threshold, calculated as the mean force plus three standard deviations (SD) from the signal during the “rest” period, and was sustained above this level for at least 100 ms (Tan et al., 2014a). Similarly, movement offset was defined as the time before the gripper force fell below this threshold and remained there for at least 100 ms. Error was calculated at each time point by determining the distance of the cursor from the ideal line (the midline of the streets) using Heron’s Formula. The total error for each street was constituted by the root mean square of errors at all time points (Schoenfeld et al., 2021).

### 2.4 MEG and MRI data acquisition

Whole-head MEG recordings were acquired in a magnetically shielded room at the Oxford Centre for Human Brain Activity (OHBA) using a 306-channel MEGIN Triux™ Neo system (MEGIN, Stockholm, Sweden). This system includes 204 planar gradiometers and 102 magnetometers. Head position was monitored using five head position indicator (HPI) coils attached behind the left and right ear lobe, above the right eyebrow, on the left and right forehead just below the hairline. The locations of the HPI coils, three anatomical fiducial points (nasion, left and right pre-auricular points) and at least 200 additional points of the head surface were digitized using a Polhemus FastTrak 3D magnetic digitization system (Polhemus, Vermont, United States) prior to MEG acquisition. Both vertical and horizontal electro-oculograms as well as electrocardiograms were recorded to monitor eye blinks, horizontal eye movements and cardiac activity, respectively. MEG data were sampled at 1000 Hz with a 0.03-330 Hz bandpass filter applied during digitization. Visual stimuli were projected to 31 x 55 cm on a screen placed 120 cm in front of the participant using a Panasonic PT D7700E projector (Panasonic, Osaka, Japan) with a refresh rate of 60 Hz.

T1-weighted structural magnetic resonance imaging (MRI) scans were acquired using a WIN Siemens 3 Tesla Prisma scanner (Siemens, Erlangen, Germany) either at the Centre for Functional Magnetic Resonance Imaging of the Brain (FMRIB) or OHBA. The protocol employed a rapid gradient echo (MPRAGE) sequence, which lasted 8 minutes. The scanning parameters were set as follows: time of repetition (TR) = 1900 ms, time of echo (TE) = 3.96 ms, voxel size = 1 x 1 x 1 mm, field of view (FOV) = 256 x 256 mm (including nose and ears), slice thickness = 1 mm, and 192 slices per slab.

### 2.5 MEG data analysis

MEG data analyses were conducted using Python 3.10. The MEG data were firstly maxfiltered using the “maxwell_filter” function from MNE-Python for noise removal, bad channel detection, and head movement correction (Gramfort et al., 2013). Subsequent preprocessing steps were performed using OHBA Software Library (OSL) (Quinn et al., 2022). The continuous data were downsampled to 250 Hz to reduce computational demands. A bandpass filter from 0.5 to 100 Hz and notch filters at 50 and 100 Hz were applied. Bad segments were identified by segmenting the data into 500 ms chunks and applying a generalized extreme Studentized deviate algorithm at a significance level of 0.1. Independent Component Analysis (ICA) based on FastICA (Hyvarinen, 1999) decomposition was employed to automatically correct stereotypical heart and eye movement artefacts by correlating the independent components with ECG and EOG channels. Next, any remaining bad channels were interpolated using spherical spline interpolation (Perrin et al., 1989). Two out of 43 participants were excluded from further analysis due to excessive artefacts.

Following preprocessing, source localization was performed to estimate the brain origins of the MEG signal. The MEG data were co-registered with individual T1-weighted MRI images for each participant using headshape points. Three participants lacked individual anatomical scans, for these participants the MNI152-T1 standard template was used. Prior to beamforming, the data were filtered into two distinct frequency bands: 5 to 45 Hz primarily for beta power and 60 to 90 Hz for gamma power. The sensor-level data were then projected onto an 8 mm grid inside the inner skull using a Linearly Constrained Minimum Variance (LCMV) beamformer (Van Veen and Buckley, 1988; Westner et al., 2022). A noise covariance matrix used to calculate the beamformer was estimated using the sensor-level data for each subject and regularised to a rank of 60 using principal component analysis (PCA). Then, the MEG data were parcellated into 52 anatomically informed regions of interest (Glasser et al., 2016; Kohl et al., 2023). Following this, symmetric orthogonalization was applied to the time courses of these parcels to minimize spatial leakage, and dipole sign flipping was used to align the signs of each parcel time course across subjects (Gohil et al., 2024).

Next, the data were epoched, aligned to movement onset (from −5 to +10 seconds relative to movement onset for analysing beta ERD and gamma ERS) and movement offset (from −2 to +5 seconds relative to movement offset exclusively for beta ERS analysis). Time-frequency transformation of the MEG data was conducted using the multitaper method, as implemented in MNE-Python. For this analysis, the number of cycles per frequency was dynamically defined to be half of the frequency and the time-bandwidth product was set at 6. Baseline correction was applied to each epoch within the interval from −1.5 to −1 seconds relative to movement onset, and the results were converted to decibel (dB) units to normalize power.

In the beta band (13 to 30 Hz), we extracted beta ERD (defined from movement onset to offset) and beta ERS (from movement offset to offset + 1 second, see Figure 2). In the gamma band (60 to 90 Hz), we extracted gamma ERS (from movement onset to onset + 0.5 seconds, see Figure 2). These analyses were specifically focused on the left and right somatosensory and motor cortex areas (Rueda-Delgado et al., 2017).

**Figure 2.**
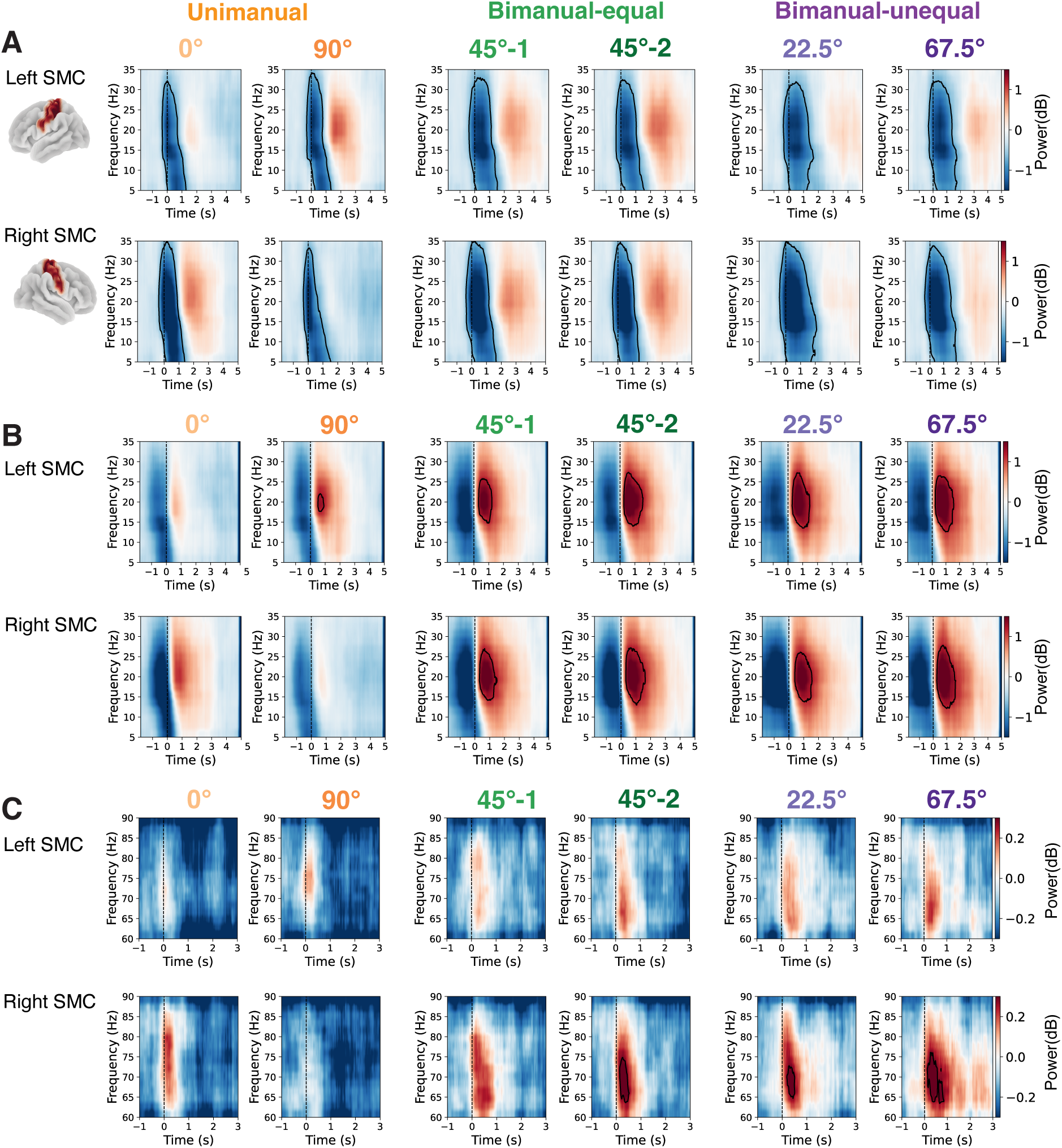
Time-frequency responses to hand movement in the sensorimotor cortex. (A) ERD predominantly in the beta (13-30 Hz) band following movement onset in both left and right sensorimotor cortices (SMC) during unimanual and bimanual conditions. Time at 0 s represents movement onset. (B) Beta ERS following movement offset, predominantly in the contralateral sensorimotor cortex during the unimanual condition, and in both sensorimotor cortices during bimanual conditions. Time at 0 s represents movement offset. (C) Gamma (60-90 Hz) ERS following movement onset, primarily in the contralateral sensorimotor cortex during the unimanual condition, and in both sensorimotor cortices during bimanual conditions. Time at 0 s represents movement onset. The two columns for the bimanual-equal condition represent the two 45° streets included in each trial. One 45° street from each trial was randomly assigned to one set of 45° streets (45°-1), and the other to a separate set of 45° streets (45°-2). Black outlines in the time-frequency plots indicate significant clusters with *p* < 0.05 by cluster-based permutation test.

For the different movement conditions (unimanual, bimanual-equal, bimanual-unequal), the principal sensorimotor cortex (P-SMC) is defined as the sensorimotor cortex (SMC) contralateral to the hand exerting more or full force, whereas the auxiliary sensorimotor cortex (A-SMC) refers to the SMC ipsilateral to the hand exerting more or full force (Figure 3A). More specifically, in the unimanual condition, the P-SMC is located in the right hemisphere for movements along the 0° street and in the left hemisphere for the 90° street. In the bimanual-equal condition, one 45° street from each trial is randomly assigned to have the right sensorimotor cortex as the P-SMC and the other to have the left sensorimotor cortex as the P-SMC. For the bimanual-unequal condition, the P-SMC is determined by the hand exerting more force, with the right sensorimotor cortex serving as the P-SMC for the 22.5° street and the left sensorimotor cortex serving as the P-SMC for the 67.5° street. Correspondingly, the A-SMC in each condition is the sensorimotor cortex on the opposite side of the P-SMC.

**Figure 3.**
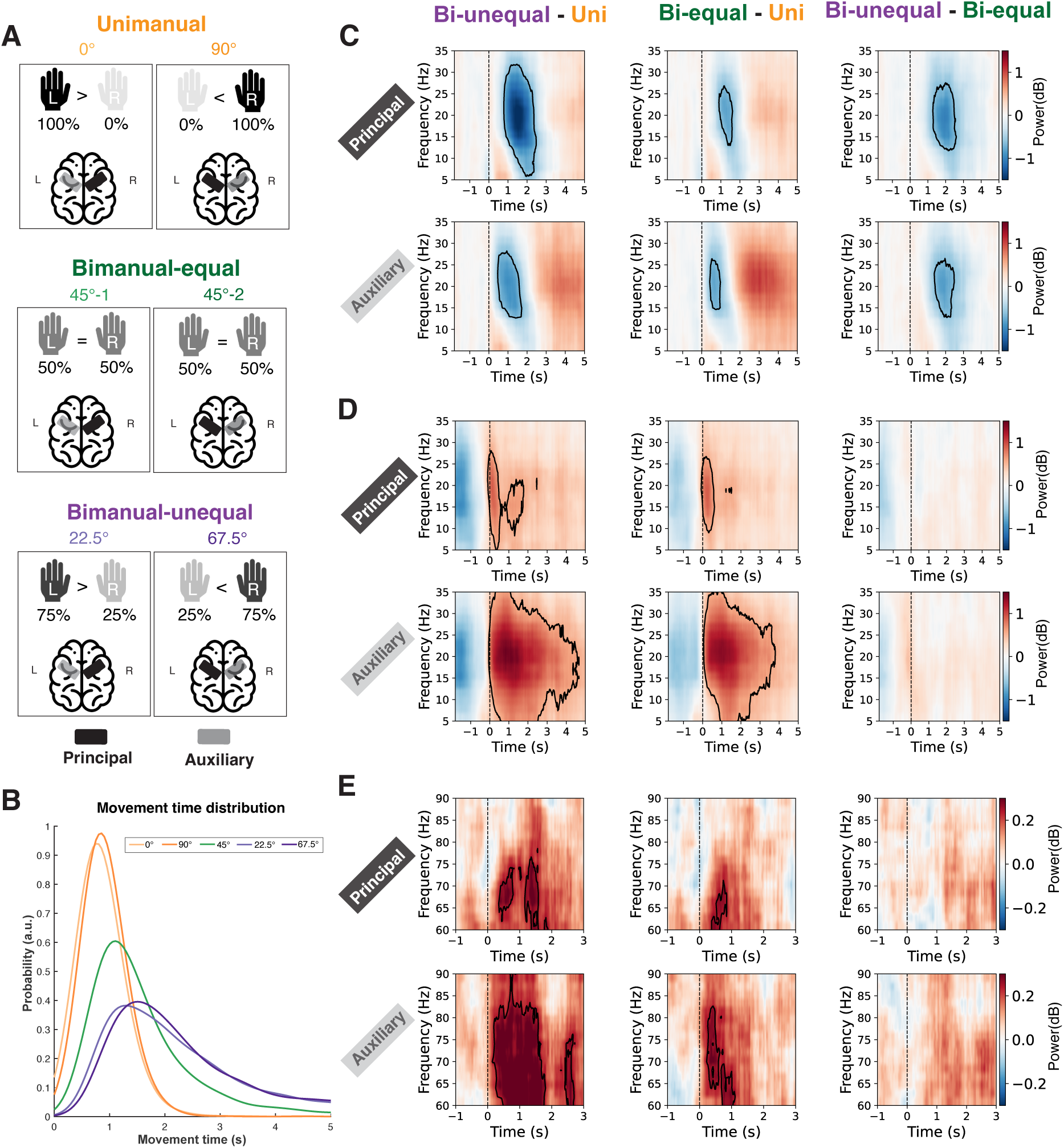
Time-frequency response differences in the sensorimotor cortex for unimanual, bimanual-equal, and bimanual-unequal conditions. (A) The black regions indicate the principal sensorimotor cortex (P-SMC), and the grey regions indicate the auxiliary sensorimotor cortex (A-SMC) for the given street. For the unimanual condition, the P-SMC is on the right hemisphere for the 0° street and on the left hemisphere for the 90° street. In the bimanual-equal condition, one 45° street from each trial was randomly assigned to 45°-1 (P-SMC in the right hemisphere) and the other to 45°-2 (P-SMC in the left hemisphere). For the bimanual-unequal condition, the P-SMC corresponds to the hand exerting more force (right hemisphere for the 22.5° street and left hemisphere for the 67.5° street). (B) Distribution of movement times across five street angles. (C) Greater beta ERD during the movement is observed in two bimanual conditions compared to the unimanual condition (left and middle panels) and in bimanual-unequal compared to bimanual-equal conditions (right panel). (D) Greater beta ERS after movement offset is observed in bimanual compared to unimanual conditions (left and middle panels), with no significant difference between bimanual-unequal and bimanual-equal conditions (right panel). (E) Greater gamma ERS after movement onset is observed in bimanual compared to unimanual conditions (left and middle panels), with no significant difference between bimanual-unequal and bimanual-equal conditions (right panel). Black outlines indicate significant clusters with *p* < 0.05 by cluster-based permutation test.

### 2.6 Statistical Analysis

The movement time and errors were compared between conditions using two-tailed paired t-tests with false discovery rate (FDR) correction (Figure 1). To assess the presence of significant ERD and ERS in the time-frequency domain, cluster-based non-parametric permutation tests were performed using 1000 sign-flipping permutations implemented in the Python toolbox OSL-dynamics (Gohil et al., 2023). These tests compared the observed power against a null hypothesis of zero within specified time windows and frequency ranges as shown in Figure 2. Similarly, cluster-based permutation tests were conducted to identify differences between movement conditions (Figure 3).

To investigate learning effects, linear mixed-effects models (LMMs) were implemented using the lme4 package in R. Prior to model fitting, the 50 trials were split into 5 blocks, each containing 10 trials, and trials within blocks were averaged per street angle. Next, the main effect of “block” was tested for each street angle, by including a fixed effect for “block” (five levels, each block corresponding to ten consecutive trials) and a random intercept for “subject”. To test if the “block” main effect differed across different street angles, a fixed effect for “street angle” (five levels: 0°, 22.5°, 45°, 67.5° and 90°) was included in the model to examine the “block” × “street angle” interaction.

Additionally, to explore the correlations between the behavioural changes (i.e., MT and error) and the neural changes (i.e., changes in beta ERD, beta ERS, and gamma ERS), further LMMs were employed. Specifically, these models included “neural change” as the dependent variable, with fixed effects for “behavioural change” (difference between blocks 1 and 5) and “street angle”, and a random intercept for “subjects” (Grigutsch et al., 2024).

## 3. Results

### 3.1 Increasing complexity of bimanual movements results in worse task performance

As would be expected the movement time (MT) for the unimanual conditions was significantly shorter compared to both bimanual-equal (0.88 ± 0.04 s vs 1.69 ± 0.12 s, mean ± SEM, t(40) = 8.07, *p* < 0.001) and bimanual-unequal conditions (0.88 ± 0.04 s vs 3.47 ± 0.35 s; t(40) = 7.80, *p* < 0.001, Figure 1D, left). Moreover, MTs were shorter for the bimanual-equal condition than the bimanual-unequal condition (t(40) = 5.91, *p* < 0.001). Similarly, movement errors were lower in unimanual condition than both the bimanual-equal condition (0.18 ± 0.05 vs 16.69 ± 0.64, t(40) = 26.16, *p* < 0.001) and the bimanual-unequal condition (0.18 ± 0.05 vs 23.58 ± 0.46, t(40) = 52.43, *p* < 0.001, Figure 1D, right). Errors in the unimanual condition almost always occurred when subjects used the incorrect hand. There were also lower errors in the bimanual-equal condition than the bimanual-unequal condition (t(40) = 12.39, *p* < 0.001).

### 3.2 Increasing complexity of movements results in greater movement-related brain activity

As expected, we observed a significant beta ERD from movement onset for approximately 1.5 seconds in both the left and right sensorimotor cortices in all conditions (Figure 2A). This was followed by significant beta ERS which persisted for approximately 1-1.5 seconds from movement offset. The beta ERS was predominantly in the contralateral sensorimotor cortex following the unimanual condition and in both sensorimotor cortices following bimanual conditions (Figure 2B). Gamma power exhibited significant ERS from movement onset for approximately 0.5-1 seconds, primarily in the contralateral sensorimotor cortex during the unimanual condition, and in both sensorimotor cortices during bimanual conditions, though it was more pronounced in the right hemisphere (Figure 2C).

We then went on to compare the movement-related changes in beta and gamma power across the task conditions. There was a significantly greater (i.e. more negative) beta ERD within both the principal and auxiliary sensorimotor cortices (Figure 3C) in both bimanual conditions compared to the unimanual condition. There was also a significantly greater ERD in the bimanual-unequal condition than the bimanual-equal condition in both SMC.

As might be expected, we observed an extended and more pronounced beta ERS within the auxiliary SMC in the bimanual conditions compared with the unimanual condition (Figure 3D). However, a significantly greater beta ERS was also observed for the bimanual conditions than the unimanual conditions in the principal SMC region, which was engaged in both the unimanual and bimanual conditions. Similarly, the gamma ERS was greater in bimanual conditions than in the unimanual condition, particularly within the auxiliary sensorimotor cortex (Figure 3E). Taken together, these findings suggest that there is increased recruitment of both the primary and auxiliary sensorimotor cortices during bimanual compared with unimanual conditions.

### 3.3 Motor learning leads to faster and more accurate movements

Next, we wanted to investigate how behaviour changed as participants performed the task. As would be expected, participants’ performance improved with practice, as demonstrated by a significant decrease in MT for each street angle individually [significant main effect of *block* for each street angle: 0° (β = −0.036, SE = 0.007, *p* < 0.001), 90° (β = −0.017, SE = 0.008, *p* = 0.024), 45° (β = −0.265, SE = 0.040, *p* < 0.001), 22.5° (β = −1.063, SE = 0.146, *p* < 0.001), and 67.5° (β = −0.710, SE = 0.104, *p* < 0.001) (Figure 4A)].

**Figure 4.**
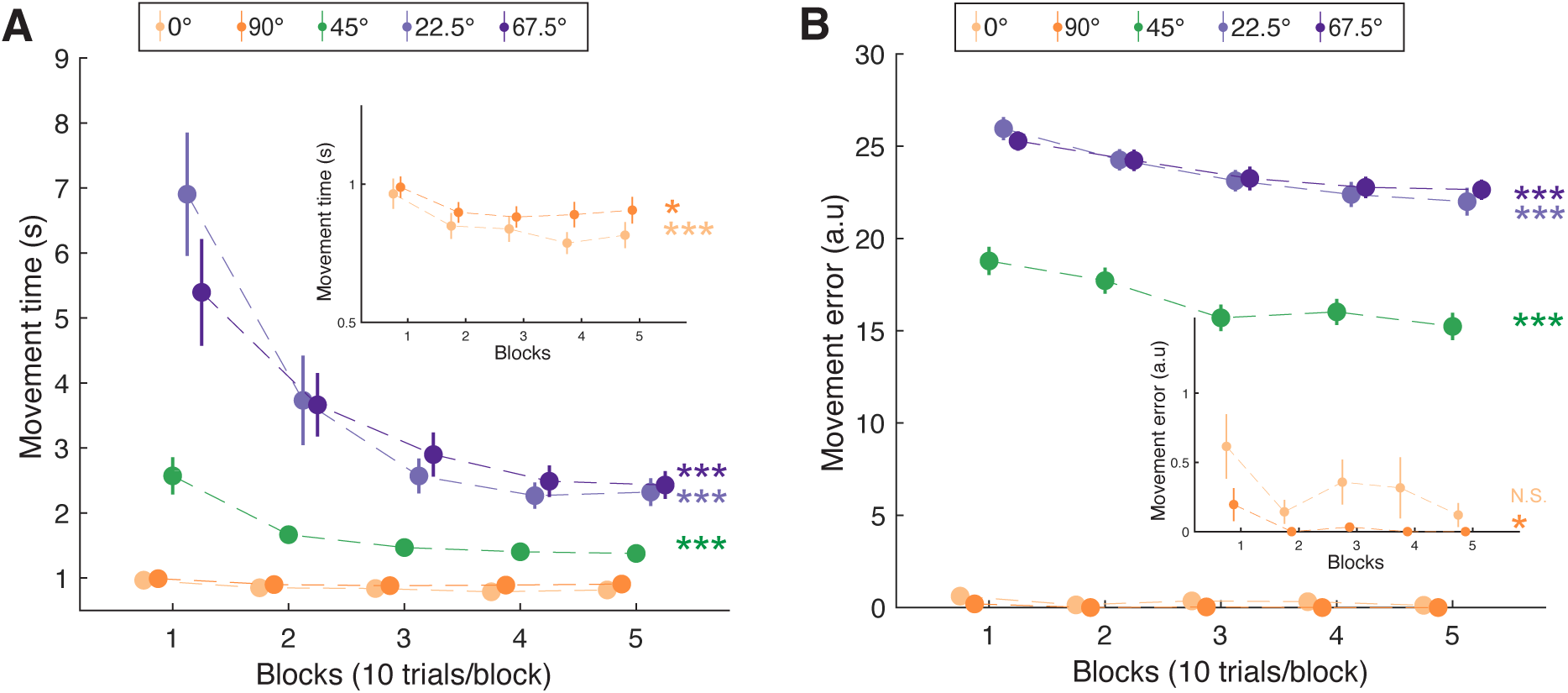
Behavioural performance across blocks for different street angles. (A) Movement time significantly decreases across blocks for all street angles. (B) Movement error shows a significant decrease across blocks for all street angles except the 0°. Each dots represent the mean performance for a block (10 trials per block) averaged across all subjects, with error bars representing the standard error of the mean across subjects. Statistical significance for the main effect of the factor *“block”* is shown as: N.S. (not significant), * *p* < 0.05, **\*\*\*** *p* < 0.001.

To investigate the influence of street angle on learning dynamics, we conducted a linear mixed model analysis on MT by further incorporating *street angle* as a fixed effect and revealed a significant interaction between *block* and *street angle* (β = −0.239, SE = 0.032, *p* < 0.001), indicating that the magnitude of movement time reduction varied as a function of *street angle*. Movement errors significantly decreased during task performance for all street angles except 0° (0°: *p* = 0.098; all other street angles: *p* < 0.05) (Figure 4B). Similarly to MT, there was also a significant *block* and *street angle* interaction for movement error (β = −0.212, SE = 0.076, *p* = 0.005).

### 3.4 Motor learning modulates beta power

Neural responses also changed during task performance. We observed a significantly weaker beta ERD (i.e. the ERD became less negative) in the left, principal SMC as participants learnt the task both for 90° (β = 0.104, SE = 0.026, *p* < 0.001) and 67.5° (β = 0.142, SE = 0.028, *p* < 0.001; Figure 5A). No significant changes were detected at 0°, 22.5°, or 45° (*p* > 0.05), and the changes in ERD across blocks did not significantly differ among street angles [*block × street angle* interaction, β = 0.017, SE = 0.010, *p* = 0.084; Figure 5A].

**Figure 5.**
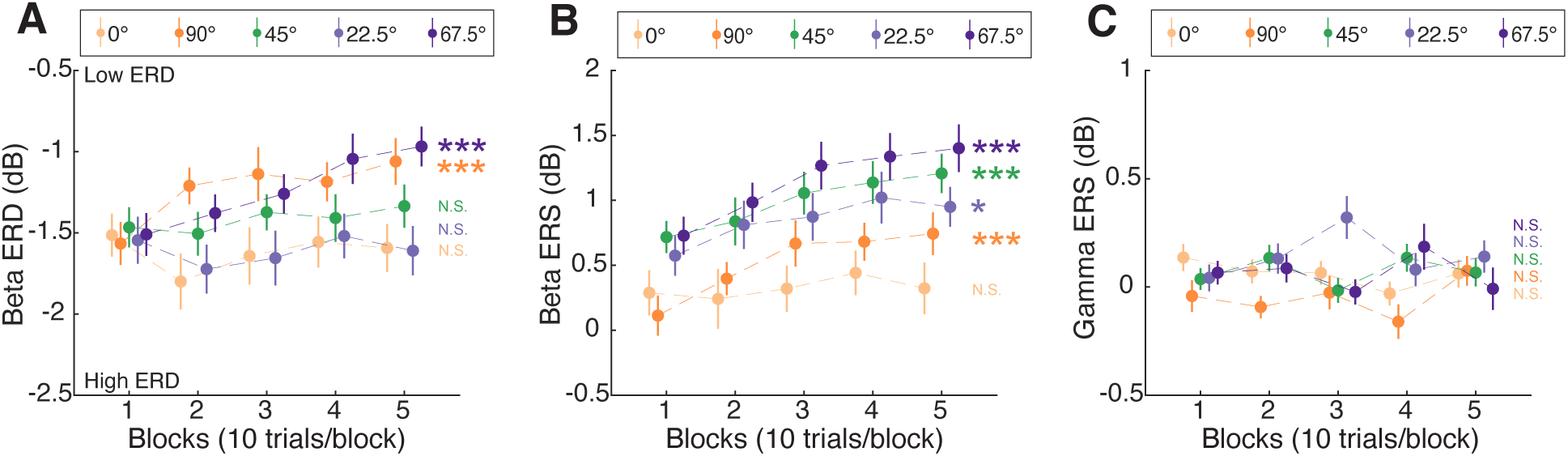
Cortical responses in the principal sensorimotor cortex across blocks for different street angles. (A) Beta ERD (from movement onset to offset) in the P-SMC as a function of blocks. Significant changes in beta ERD were observed across blocks for the 90° and 67.5° streets. (B) Beta ERS (from movement offset to 1 s after) in the P-SMC as a function of blocks. Beta ERS significantly increased across blocks for all street angles except the 0° street. (C) Gamma ERS (from movement onset to 0.5 s after) in the P-SMC as a function of blocks. No significant changes were detected in any street angle. Data points represent the mean power for a block (10 trials per block) averaged across all subjects, and error bars represent the standard error of the mean across subjects. Statistical significance for the main effect of the factor “block” is indicated as: N.S. (not significant), * p < 0.05, **\*\*\*** p < 0.001.

Beta ERS in the principal SMC significantly increased with task performance in 90° (β = 0.155, SE = 0.033, p < 0.001), 45° (β = 0.128, SE = 0.032, p < 0.001), 22.5° (β = 0.096, SE = 0.037, *p* = 0.010), and 67.5° (β = 0.170, SE = 0.036, *p* < 0.001) streets, with no significant change in 0° streets (β = 0.027, SE = 0.042, *p* = 0.525; Figure 5B). There was no significant *block* × *street angle* interaction (β = 0.023, SE = 0.012, *p* = 0.080).

No significant block effects were detected in gamma ERS across any conditions in the P-SMC (p > 0.05, Fig. 5C). Further analysis of neural responses in A-SMC yielded a similar pattern of results to those observed in the P-SMC (Supplementary Figure S1).

### 3.5 Correlation between changes in task performance and neural responses

Finally, we examined the relationships between the learning-related changes in behaviour and changes in neural responses (Block-5 minus Block-1) by using linear mixed models. Given that the unimanual streets frequently contained no errors, our correlation analysis primarily focused on the bimanual conditions, though the overall results remained consistent when including all movement conditions, as detailed in Supplementary Figure S3.

This analysis revealed that learning-related changes in MT showed a trend toward significant correlation with changes in beta ERD in P-SMC (β = 0.036, *p* = 0.045; with all conditions included: β = 0.030, *p* = 0.089, see Supplementary Figure S3A), indicating that reductions in MT were associated with increasingly negative beta ERD. However, no such association was observed between movement error and beta ERD (β = −0.007, *p* = 0.724) (Figure 6A). Changes in MT did not correlate with changes in beta ERS (β = −0.014, *p* = 0.456). However, a greater reduction in movement errors was associated with an increase in beta ERS (β = −0.049, *p* = 0.018), indicating that improvements in task accuracy, reflected by decreased errors, were correlated with increased beta ERS (Figure 6B). Changes in gamma ERS were not related to changes in either MT or movement error (Figure 6C).

**Figure 6.**
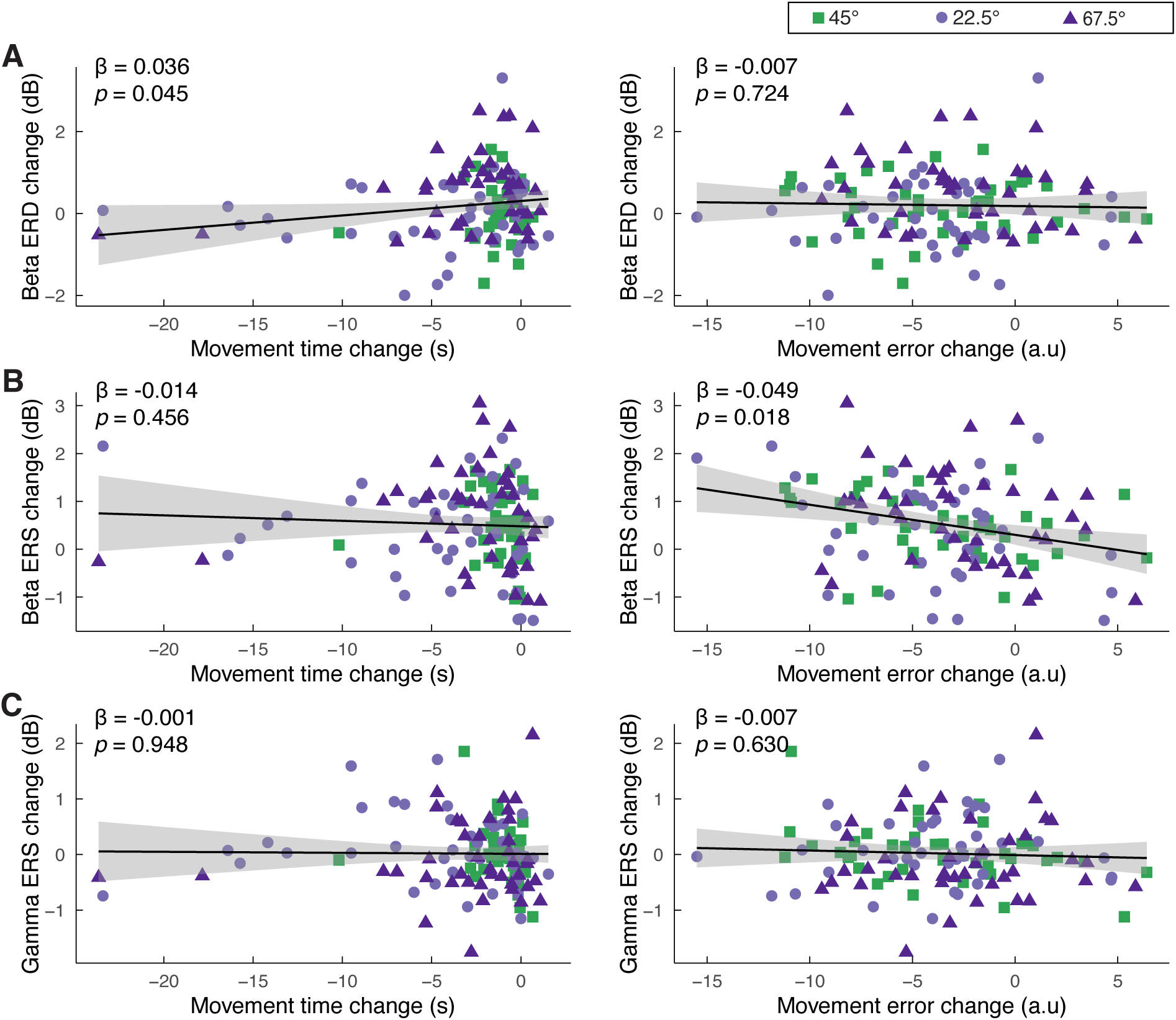
Relationship between learning-related changes in behaviour and neural responses. Linear mixed model analysis was used to assess the association between changes in movement performance and changes in neural responses in P-SMC. (A-C) show the changes in beta ERD, beta ERS, and gamma ERS as a function of movement time change (left) and movement error change (right). (A) Beta ERD changes exhibit a significant positive relationship with movement time change. No significant relationship is observed with movement error change. (B) Beta ERS changes reveal a significant negative relationship with movement error change. No significant relationship is found with movement time change. (C) Gamma ERS changes show no significant relationships with either movement time or movement error changes. Data points represent individual trials for the 45°, 22.5°, and 67.5° streets, coloured in green, light purple, and dark purple, respectively. The solid black lines depict the fitted regression lines from the linear mixed models, with shaded areas representing the 95% confidence intervals.

### 3.6 Left and right SMC show differing beta and gamma power

We observed a significant difference in neural responses between left and right SMC. Beta ERD was larger in the right SMC compared to the left SMC, when comparing 0° (right PMC as the P-SMC) and 90° (left PMC as the P-SMC) (β = −0.389, SE = 0.060, *p* < 0.001), and 22.5° (right PMC as the P-SMC) and 67.5° (left PMC as the P-SMC) (β = −0.379, SE = 0.021, *p* < 0.001). The gamma ERS showed a similar hemispheric dominance to the beta ERD, with greater power in the right compared with the left SMC (Figure 5). This right SMC dominance was also seen in the bimanual-equal condition, where the right SMC exhibits greater beta ERD and larger gamma ERS compared to the left SMC (Supplementary Figure S2).

## 4. Discussion

In this study, we investigated the behavioural and neural changes during performance and learning of a novel bimanual task with different levels of bimanual interaction. As hypothesised, unimanual movements were associated with shorter movement times and higher accuracy than bimanual movements. Among the bimanual movements, the unequal movements, which had a higher degree of complexity, proved more challenging as expected, exhibiting longer movement times and lower accuracy than the bimanual movements which required equal contributions from each hand. On the neural level, as expected, we observed significant movement-related activity in terms of beta ERD, beta ERS, and gamma ERS across all movement conditions, with bimanual movements eliciting enhanced responses.

As seen previously (Schoenfeld et al., 2021), as participants performed the task their performance improved in almost all movement conditions, reflected by a decrease in movement times and errors. These changes suggest that participants were not merely repeating movements, but rather that practice refined their motor skills. As hypothesised, in parallel with this improved behaviour, we observed an increase in beta ERS during learning, which correlated with the reduction in errors in the task. Contrary to our hypothesis, we observed lower power in beta ERD (i.e. a smaller deflection) during learning in conditions where the principal SMC was in the left hemisphere.

### 4.1 Neural dynamics are modulated during movement

Consistent with previous findings, we observed that beta ERD occurs during movement execution within the 13-30 Hz frequency range. This reduction in beta power is thought to reflect the release of cortical inhibition and activation of the motor network (Pfurtscheller et al., 1997a; Tinkhauser et al., 2019). Beta ERD is frequently reported bilaterally in electrophysiological (EEG) studies even during unimanual movements. This contrasts with the more pronounced lateralization seen in hemodynamic responses recorded simultaneously with fMRI-EEG. However, further functional MRI studies indicate that ipsilateral activity during unimanual movements likely results from an efference copy of motor signals generated in the contralateral cortex. This activity aids in supressing mirror-symmetric actions and ensures coordinated motor control during unimanual tasks, preventing unintended movements on the opposite side (Ames and Churchland, 2019; Diedrichsen et al., 2013).

Post-movement beta ERS occurs after movement termination and is believed to reflect a prolonged period of increased corticomuscular coherence following voluntary movements (Feige et al., 2000). This increase in beta power is also suggested to be associated with motor plasticity and skill retention (Ghilardi et al., 2021; Peter et al., 2022). Our findings align with previous magnetoencephalography and electrophysiological studies, which report a bilateral beta ERS, which is greater in the sensorimotor cortex contralateral to movement (Devos et al., 2006; Jurkiewicz et al., 2006).

We observed gamma activity in the 60-90 Hz range coinciding with movement onset and lasting approximately 0.5-1 seconds, which demonstrates strong lateralization to the SMC contralateral to movement. This unilateral pattern and relatively short time window supports the view that gamma ERS has greater spatial and temporal specificity compared to beta activity (Crone et al., 1998; Muthukumaraswamy, 2010). The strong lateralization supports the hypothesis that gamma ERS is crucial for motor coordination and initiation, particularly in encoding contralateral movements (Miller et al., 2009; Mooshagian et al., 2021).

Finally, we observed a right hemisphere dominance for beta ERD and gamma ERS, which aligns with findings of larger beta ERD in the right sensorimotor cortex contralateral to the non-dominant hand in both unimanual and bimanual tasks (Houweling et al., 2008). Additionally, studies on unimanual movements show larger activity changes in the contralateral sensorimotor cortex when moving the left finger compared to the right (Jäncke et al., 1998), supporting hemispheric asymmetries in motor control.

### 4.2 Greater task complexity leads to greater movement-related activity

Bimanual movements are inherently more demanding than unimanual movements, due to the increased need for temporal coordination between the hands (Serrien and Brown, 2002), which in our task is especially apparent in the bimanual unequal movements. In line with that, we demonstrated that beta ERD, beta ERS, and gamma ERS are all more pronounced in bimanual movements compared to unimanual movements.

These observations are consistent with recent non-human primate studies using local field potential recordings from the parietal reach region, which demonstrated task-specific variations in beta power across different bimanual and unimanual tasks (Mooshagian et al., 2021). This study suggested that beta power in both cortices encodes movement-specific information for both hands. In contrast, gamma power likely only encodes contralateral arm movements, offering limited insight into ipsilateral arm actions (Mooshagian et al., 2021).

Correspondingly, our analysis revealed that beta ERD varies across different motor tasks in both hemispheres, supporting its role in encoding task complexity. However, gamma ERS was significantly activated in the A-SMC during bimanual tasks, but not in unimanual movements, resulting in a pronounced difference primarily in the A-SMC between bimanual and unimanual movements. Collectively, these findings illustrate that beta power provides a rich, multifaceted view of motor coordination involving both hands, but gamma power is more narrowly focused on contralateral movements.

### 4.3 Increases in beta ERS correlate with error reduction during learning

We demonstrated significant increases in post-movement beta ERS across the task, except at the 0° street angle, supporting the hypothesised relationship between increases in learning-related beta ERS and error reduction. Previous studies have demonstrated that post-movement beta ERS in the sensorimotor cortex is associated with learning processes (Tan et al., 2016, 2014b; Torrecillos et al., 2015). However, the precise functional role of beta ERS remains unclear.

Early theories suggest that beta ERS represents a return to an idling state of the sensorimotor cortex, indicative of reduced sensorimotor cortical excitability. According to these theories, beta ERS serves an active inhibitory function by “clearing out” the motor program after movement execution and resetting the neural network to its baseline state (Kilavik et al., 2013; Schmidt et al., 2019). Recent studies, however, propose that the role of beta ERS extends beyond simply reflecting an inactive or inhibited cortical state. Beta ERS has been shown to relate to movement error (Tan et al., 2016, 2014a; Torrecillos et al., 2015). Consistent with this, our finding that the learning-related increase in beta ERS correlates with learning-related decrease in error, suggests that beta ERS may be involved in the comparison between planned and executed movements. Additional research suggests that enhancements in beta ERS are linked to rule acquisition, stabilizing and amplifying the post-decision “beta-state” to strengthen neural representations of newly learned rules (Bogaerts et al., 2020; Pavlova et al., 2023). These modifications may reflect changes in higher-order cognitive processes rather than mere improvements in movement parameters.

### 4.4 Beta ERD is reduced during learning, but only in the left, principal SMC

Unlike beta ERS, we did not observe consistent changes in beta ERD during practice across different movement conditions: only conditions in which the left sensorimotor cortex was principally involved showed a less negative deflection in ERD. The role of beta ERD in learning is not consistent across the previous literature. For example, one study reports a practice-induced strengthening in beta ERD (i.e. a more negative deflection) (Zich et al., 2018), indicating enhanced neural efficiency with practice. In contrast, other studies find no significant differences in sensorimotor beta ERD during learning (Pavlova et al., 2023; van der Cruijsen et al., 2021), challenging the notion that beta ERD reflects learning-related neural adaptations.

All our participants were right-handed, hence the left SMC is in the dominant hemisphere in all participants. It is not possible, therefore, to unpick whether the differences between the pattern of learning-related changes in the left and right SMCs are due to intrinsic hemispheric differences or rather reflect the dominant hemisphere.

### 4.5 Gamma ERS does not change with learning

Our study observed no significant changes in gamma ERS during practice in any movement conditions, suggesting that gamma ERS may not play a critical role in motor learning quantified by reduced movement time and reduced error. This observation appears to contradict the view that gamma activity plays a prokinetic role in movement, and motor skill acquisition (Muthuraman et al., 2020; Zich et al., 2021). For instance, studies using 75 Hz or theta-gamma coupling transcranial alternating current stimulation (tACS) have shown relationships between 75Hz activity and skill acquisition (Akkad et al., 2021; Nowak et al., 2018). However, they studied thumb abduction acceleration rather than the movement time and error used here; it may therefore be that 75Hz activity more directly reflects force modulation than other aspects of motor performance. Interestingly, it has been proposed that the efficiency of gamma tACS may occur through the modulation of beta activity rather than gamma activity (Sugata et al., 2018), which might at least partially explain the pattern of results observed here.

Finally, it has been hypothesised that early changes in GABA inhibition after tACS are linked with motor learning (Nowak et al., 2018), and gamma ERS typically peaks during early skill acquisition (Madhavan et al., 2019; Perfetti et al., 2011). This is in line with the finding that changes in gamma ERS occur mainly in the initial phases of learning (Haverland et al., 2024), to which the design of our task may be relatively insensitive.

### 4.6 Task considerations

Reflecting the different task demands of different conditions, movement times and errors differed across task conditions. Additionally, as participants learnt our task, their movement times and errors decreased. The differences in neural responses we observe between conditions and across the task might therefore result from variations in movement times, rather than differences in bimanual co-ordination per se, such that task conditions with longer movement times might exhibit more pronounced neural changes due to prolonged neural activity. However, the temporal specificity of significant differences in neural dynamics makes this explanation less likely. For example, the gamma ERS clusters (Figure 3E) align closely with movement onset. If the differences in gamma ERS across movement conditions were caused by the extended movement time, we would expect the cluster to appear later, during the prolonged movement phase, rather than at the movement onset when both unimanual and bimanual movements share a similar duration. This temporal specificity is in contrast with the variability and delayed onset of significant beta ERD clusters when comparing different task conditions, indicating that movement time may influence beta ERD (Figure 3C). Furthermore, correlation analyses reveal a correlation between changes in movement time and changes in beta ERD but not with changes in gamma or beta ERS, supporting the notion that the observed differences in beta ERD, but not beta ERS or gamma ERS, between conditions could be affected by movement time.

## 5. Conclusions

This study enhances our understanding of the neural mechanisms distinct to unimanual and bimanual movements, revealing differential responses of beta and gamma activities to these varying task complexities. Both activities are notably more active in bimanual movements, underscoring the increased demands for neural coordination. Furthermore, our findings advance insights into the neural underpinnings of motor learning, particularly emphasizing the functional significance of beta ERS. This suggests that specific neural rhythms can be strategically targeted to optimize motor learning and recovery processes.

## Author contributions

MW: conceptualisation, methodology, formal analysis, writing – original draft, writing – review and editing. MS: conceptualisation, investigation, writing – review and editing. CL: conceptualisation, investigation, writing – review and editing. SB: methodology, investigation, writing – review and editing. CZ: conceptualisation, methodology, supervision, and writing review and editing. CJS: conceptualisation, methodology, resources, funding acquisition, supervision, and writing – review and editing.

## Acknowledgments

We thank Ipsita Sarkar and Patricia Cambalova for their help with participant recruitment and data collection.

## Data availability

The datasets and code will be shared via the MRC BNDU data sharing platform on https://data.mrc.ox.ac.uk/.

## Rights retention text

This research was funded by the Wellcome Trust [203139/Z/16/Z]. For the purpose of open access, the author has applied a CC BY public copyright licence to any Author Accepted Manuscript version arising from this submission.

