## Supplemental Figs1-3 for "Differential beta and gamma activity modulation during unimanual and bimanual motor learning"

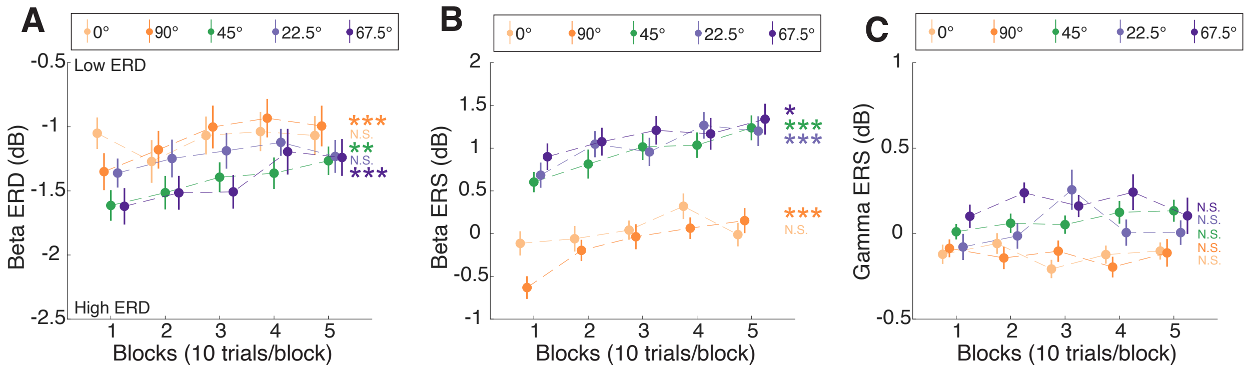


**Figure S1. Cortical responses in the auxiliary sensorimotor cortices across blocks for different street angles.** (A) Beta ERD (from movement onset to offset) as a function of blocks. Significant changes in beta ERD were observed across blocks for the 45°, 67.5°, and 90° streets. (B) Beta ERS (from movement offset to 1 s after) as a function of blocks. Beta ERS significantly increased across blocks for all street angles except the 0° street. (C) Gamma ERS (from movement onset to 0.5 s after) as a function of blocks. No significant changes were detected in any street angle.

Data points represent the mean power for a block (10 trials per block) averaged across all subjects, and error bars represent the standard error of the mean across subjects. Statistical significance for the main effect of the factor "block" is indicated as: N.S. (not significant), * $p < 0.05$, *** $p < 0.001$.

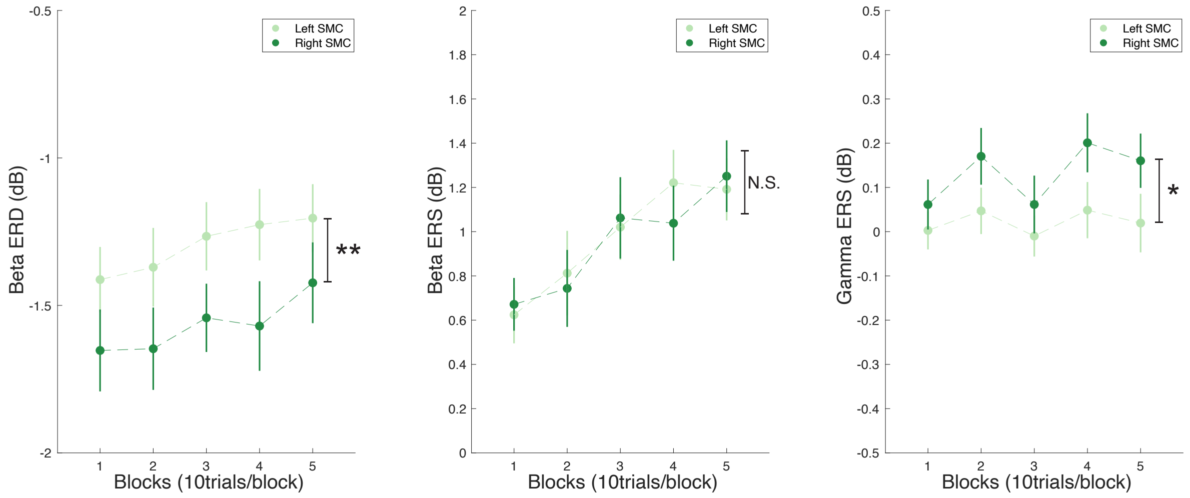


**Figure S2. Differences in time-frequency responses between the right and left sensorimotor cortex for bimanual-equal (45°) condition.** (A) Greater beta ERD is observed in the right sensorimotor cortex compared to the left sensorimotor cortex. (B) No significant difference in beta ERS after movement offset is observed between the left and right sensorimotor cortices. (C) Greater gamma ERS is observed in the right sensorimotor cortex compared to the left sensorimotor cortex.

Asterisks indicate significant differences in responses between the left and right sensorimotor cortices (* $p < 0.05$, ** $p < 0.01$). N.S. denotes not significant.

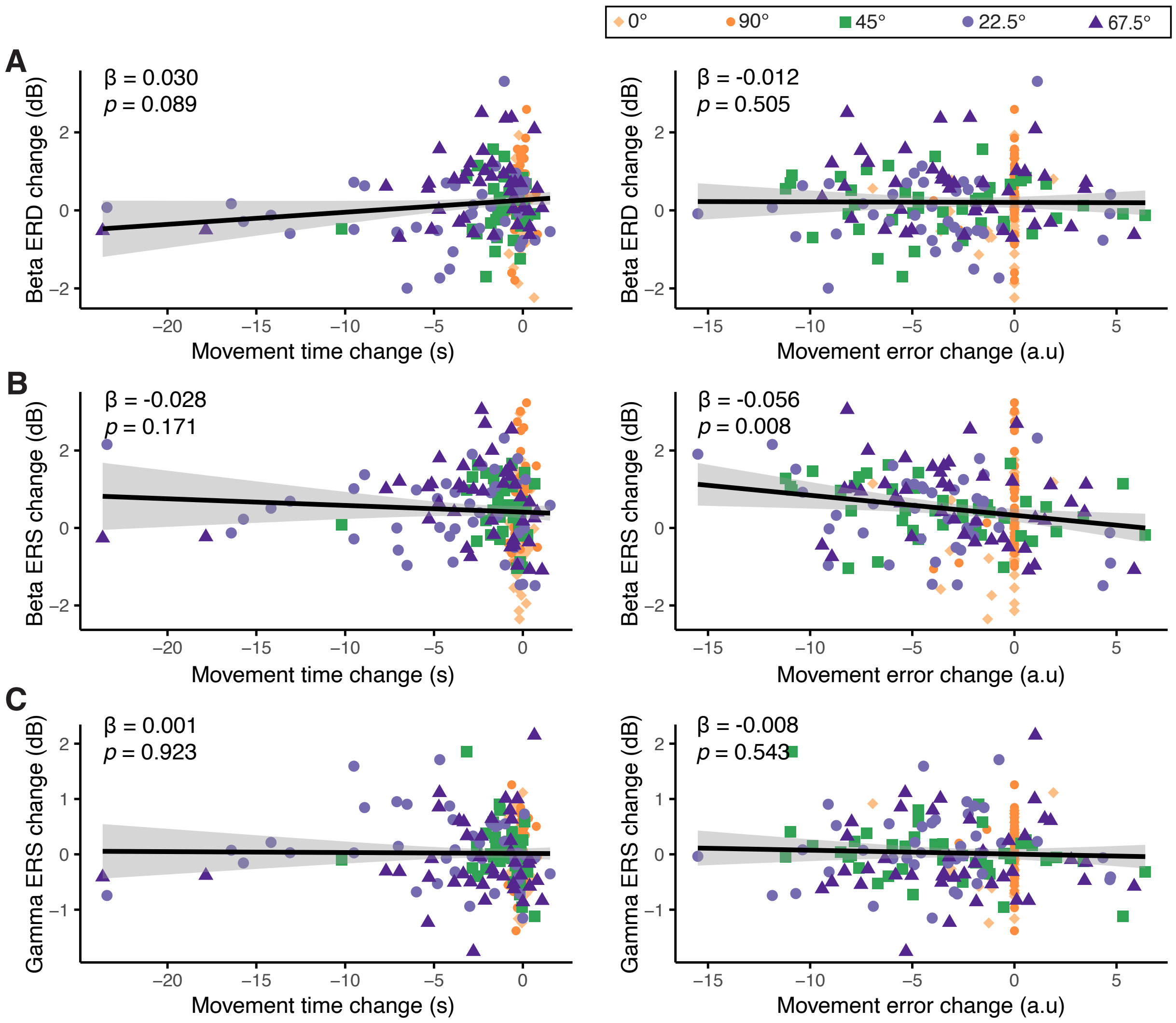


**Figure S3. Comprehensive relationship between movement performance changes and cortical responses across all street angles.** Similar to Figure 6, but includes data from all street angles (0°, 90°, 45°, 22.5°, and 67.5°). (A) Beta ERD change shows a trend of positive correlation with movement time change. (B) Beta ERS change shows a significant negative correlation with movement error change. (C) Gamma ERS change does not show significant correlations with either movement time or error changes.

Data points represent individual trials for the 0°, 90°, 45°, 22.5°, and 67.5° streets, coloured in light orange, dark orange, green, light purple, and dark purple, respectively. The solid black lines depict the regression lines from the linear mixed models, with shaded areas representing the 95% confidence intervals.
